# MatrixQCvis: shiny-based interactive data quality exploration for omics data

**DOI:** 10.1101/2021.06.17.448827

**Authors:** Thomas Naake, Wolfgang Huber

**Affiliations:** Genome Biology Unit, European Molecular Biology Laboratory, Heidelberg, 69117, Germany

## Abstract

**Motivation:** First-line data quality assessment and exploratory data analysis are integral parts of any data analysis workflow. In high-throughput quantitative omics experiments (e.g. transcriptomics, proteomics, metabolomics), after initial processing, the data are typically presented as a matrix of numbers (feature IDs × samples). Efficient and standardized data-quality metrics calculation and visualization are key to track the within-experiment quality of these rectangular data types and to guarantee for high-quality data sets and subsequent biological question-driven inference.

**Results:** We present `MatrixQCvis`, which provides interactive visualization of data quality metrics at the per-sample and per-feature level using `R’s shiny` framework. It provides efficient and standardized ways to analyze data quality of quantitative omics data types that come in a matrix-like format (features IDs × samples). `MatrixQCvis` builds upon the Bioconductor `SummarizedExperiment` S4 class and thus facilitates the integration into existing workflows.

**Availability:** `MatrixQCVis` is implemented in R. It is available via Bioconductor and released under the GPL v3.0 license.

**Supplementary information:** Supplementary Information is available at *bioRxiv* online.

## 1 Introduction

Initial first-line data quality assessment is an integral part of data analysis for subsequent biological question-driven inference. To ensure facile exploration of data quality we developed `MatrixQCVis`, implemented in the R programming language. `MatrixQCVis` provides shiny-based interactive visualization and quantification of data quality metrics at the per-sample and per-feature level. It is broadly applicable to quantitative omics data types that come in a matrix-like format (features x samples). It enables the detection of low-quality samples, outliers, drifts, and batch effects in data sets. Visualizations include boxplots and violin plots of the (count or intensity) values, mean vs standard deviation plots, MA plots (see Figure 1 a) and Hoeffding’s D statistic (non-parametric measure of independence between M and A), empirical cumulative distribution function (ECDF) plots, visualizations of the distances between samples, and multiple types of dimension reduction plots. Furthermore, `MatrixQCVis` facilitates differential expression analysis based on the `limma` (moderated t-tests, Ritchie *et al*., 2015) and `proDA` (Wald tests, Ahlmann-Eltze and Anders, 2019) packages.

**Figure 1.**
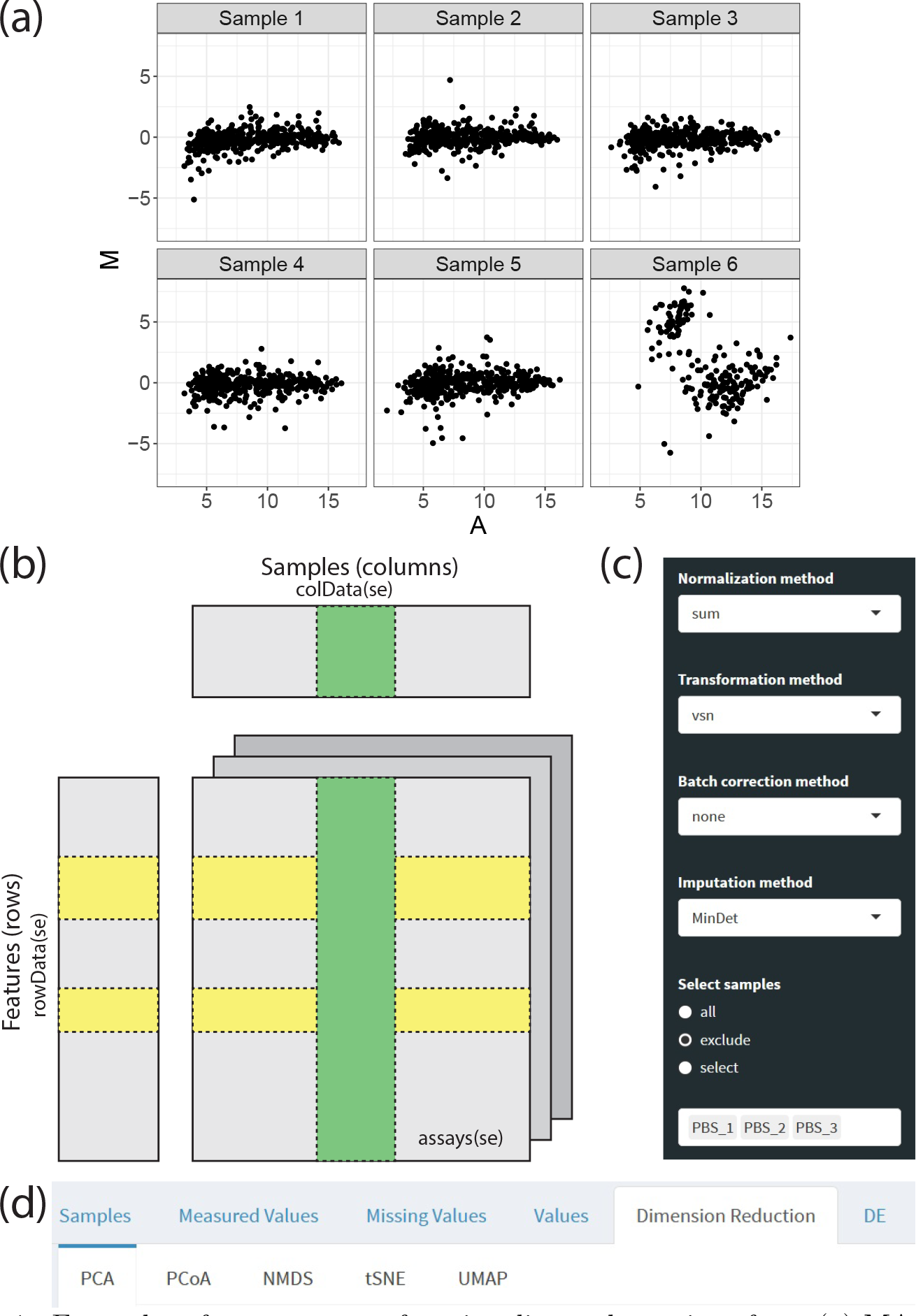
Examples of `MatrixQCVis` functionality and user interface. (a) MA plot of human plasma proteomics samples identifying a dependence between M and A values for Sample 6 indicating problems with its data. More visualizations using the clinical data sets of Jiang *et al*. (2019) and Brueffer *et al*. (2018) are shown in the Supplementary Information. (b) `MatrixQCVis` builds upon the `SummarizedExperiment` S4 class, a container for assay data (e.g. proteomics intensity values) and associated meta data on the features and samples. The figure is adjusted from the vignette of the `SummarizedExperiment` package. (c) Sidebar panel. `MatrixQCVis` enables to interactively normalize, transform, perform batch correction and impute the data set. Furthermore, samples can be excluded or selected. In the shown example, phosphate-buffered saline (PBS) samples are excluded. (d) Main panel. Navigation within `MatrixQCVis` is realized by browsing through tabs. Each visualization is embedded within a dedicated tab.

Similarly to the `iSEE` package (Rue-Albrecht *et al*., 2018), `MatrixQCVis` builds upon the widely used Bioconductor `SummarizedExperiment` S4 class (see Figure 1 b) and thus facilitates the integration into existing workflows. Compared to `iSEE`, which provides a general interface for exploring data in a `SummarizedExperiment` object, `MatrixQCVis` focuses on the upstream, initial first-line, data quality control steps and incorporates dedicated visualization capabilites for assessing data quality, although overlaps exist (e.g. dimension reduction plots). Several software packages exist that center around the assessment of data quality: among others, `arrayQualityMetrics` (Kauffmann *et al*., 2009), initially developed more than 10 years ago for the quality assessment of microarrays, that creates automatic reports of the data quality. Contrary to `arrayQualityMetrics`, which is built upon outdated visualization libraries, `MatrixQCVis` uses the `shiny` (Chang *et al*., 2021) framework and provides a high number of interactive visualizations to explore the data. `MeTaQuaC` (Kuhring *et al*., 2020), a recently published R package dedicated to metabolomics data analysis, has a special focus on the Biocrates data output and creates a static quality control report. On the other hand, `MatrixQCVis` is not restricted to a vendor-specific output and centers around interactive exploration of data quality.

We highlight the usability and functionality of the `MatrixQCVis` package in applications of clinical proteomics and transcriptomics studies in the Supplementary Information, using the data sets of Jiang *et al*. (2019) and Brueffer *et al*. (2018).

## 2 Usage scenario and user interface

Many tools and software packages exist that directly process raw data and translate these to biological discovery, but offer limited capabilities for data quality control. However, data quality control is a necessary step in omics data processing to identify samples with poor intensities and low signal-to-noise ratio, biases and outliers, batch and confounding effects, or drifts in calibration. Quality control involves the monitoring of data processing steps from raw data to processed/completed data sets (including the steps of sample normalization, batch correction, transformation, and handling of missing values) and the simultaneous assessment of quality metrics. Concomitantly, a data-quality centered workflow would facilitate consistent processing to minimize technical variance and interference. Additionally, such a workflow would ideally identify systematic trends and deviations (systematic errors) and remove these prior to data interpretation to enable sound biological information retrieval. To offer a tool to the community that addresses this gap, `MatrixQCVis` was developed.

Since `MatrixQCVis` is based on `shiny` (see Figure 1 c and d), it requires little programmatic interaction to start the quality assessment and is, thus, suitable for anyone who would like to analyze data quality in a fast, efficient, and standardized manner. `MatrixQCVis` enables users to explore data quality of rectangular data sets that are represented as a `SummarizedExperiment` object (see Figure 1 b) interactively and reproducibly. The interface to the `shiny` interface is initiated with a single call to the `shinyQC` function. A `SummarizedExperiment` object can be passed to the `shinyQC` call directly or loaded via an upload interface. Using domain-and study-specific knowledge, `MatrixQCVis` guides the user to downsample data sets on the per-sample and per-feature level.

For data entities where missing values are present in the assay structure the interface will load specific interfaces to assess the data quality with respect to missing values, e.g. tab panels to visualize the number of missing values across samples or sets of missing values across sample types, and a user interface to control imputation.

Besides the interactive quality exploration, `MatrixQCVis` enables users to download all plots and to generate markdown/HTML reports from within the `shiny` interface.

## 3 Conclusion

The `shiny` application `MatrixQCVis` generates interactive data quality workflows and facilitates to monitor data quality along the major data processing steps via several commonly applied data quality metrics and visualizations. It enables users to create a dynamic, easy-to-share and easy-to-store report using user-specified settings. `MatrixQCVis` can be integrated into existing workflows and provides a means to scrutinize the data quality of rectangular data sets in a fast, efficient, and standardized manner.

## Acknowledgements

We acknowledge financial support from the Bundesministerium für Bildung und Forschung (grant agreement no. 161L0212E). We acknowledge feedback from the SMART-CARE consortium on usability of `MatrixQCVis` and all developers and maintainers of the R/Bioconductor packages `MatrixQCVis` is built upon.

## Notes

### Competing Interest Statement

The authors have declared no competing interest.

