## Supplementary information for "MatrixQCvis: shiny-based interactive data quality exploration for omics data"

Thomas Naake and Wolfgang Huber*

The Supplementary Information showcases the functionality of the `MatrixQCvis` package by presenting an exemplary analysis workflow. While the `MatricQCvis` package entails an interactive shiny application, including also interactive plots based on the `plotly` framework, static plots are presented here.

The analysis here focuses on the clinical data sets of Jiang et al. [2019] and of Brueffer et al. [2018]. The data set of Jiang et al. [2019] is a label-free mass spectrometry proteomics data set of clinical early-stage hepatocellular carcinoma, containing in total 248 samples (124 tumor and 124 paired non-tumor tissue samples) taken prior to chemotherapy or radiotherapy. The data set of Brueffer et al. [2018] was acquired by Illumina paired-end RNA-sequencing of breast cancer biopsies and entails 405 samples.

#### 1 Preparation of the environment

Load the `MatrixQCvis` package before starting the analysis.

```
library("MatrixQCvis")
```

*European Molecular Biology Laboratory, Meyerhofstrasse 1, 69117 Heidelberg, Germany

### 2 Jiang et al. (2019): Proteomics identifies new therapeutic targets of early-stage hepatocellular carcinoma

In their study, Jiang et al. [2019] analyzed primary tumor tissues and paired non-tumor liver tissues from 110 early-stage cases of hepatocellular carcinoma (according to the Barcelona Clinic Liver Cancer stages 0 and A) related to hepatitis B virus infection. The samples were taken prior to chemotherapy or radiotherapy. The samples were analyzed using a label-free proteomics mass spectrometry approach. The analysis of Jiang et al. [2019] resulted in the stratification into the subtypes S-I, S-II, and S-III, each associated with a specific clinical outcome.

The data set was downloaded from the PRIDE database (accession number PXD006512, available at www.ebi.ac.uk/pride/archive).

Since `MatrixQCvis` requires a `SummarizedExperiment` object, we will create in the following such an object from the files containing the intensity values and the metadata of the samples.

```
## read the file with protein intensities
f <- "Jiang_et_al_2019/proteinGroups.txt"
jiang2019 <- read.delim(file = f)
rownames(jiang2019) <- jiang2019[, "Protein.IDs"]

## remove the row entries with reversed sequences
jiang2019 <- jiang2019[-grep(rownames(jiang2019), pattern = "^REV_"), ]

## retrieve the columns that contain IBAQ-processed values
jiang2019_ibaq <- jiang2019[, grep(colnames(jiang2019),
    pattern = "iBAQ[.]")]
library(stringr)
colnames(jiang2019_ibaq) <- str_remove(colnames(jiang2019_ibaq),
    pattern = "iBAQ.")
jiang2019_ibaq <- as.matrix(jiang2019_ibaq)
mode(jiang2019_ibaq) <- "numeric"

## create the object jiang2019_ibaq_na and encode the 0s by NA
jiang2019_ibaq_na <- jiang2019_ibaq
jiang2019_ibaq_na[jiang2019_ibaq_na == 0] <- NA

## create rowData
rD <- jiang2019[, c("Protein.IDs", "Majority.protein.IDs", "Protein.names",
                    "Gene.names", "Number.of.proteins", "Peptides")]
rownames(rD) <- jiang2019[, "Protein.IDs"]

## create colData
library(xlsx)
f <- "Jiang_et_al_2019/Supplementary materials for MaxQuant searching.xlsx"
```

```
cD <- read.xlsx(file = f, sheetIndex = 1)
cD <- cD[-nrow(cD), ]
rownames(cD) <- cD[, "Experiment"]
cell_type <- character(nrow(cD))
cell_type[grep(cD$Experiment, pattern = "[0-9]T$")] <- "T"
cell_type[grep(cD$Experiment, pattern = "[0-9]P$")] <- "P"
cD <- dplyr::mutate(cD, name = Experiment, cell_type = cell_type,
      patient = str_remove(cD$Experiment, pattern = "P|T"))

se <- SummarizedExperiment(assays = list(jiang2019_ibaq), colData = cD,
    rowData = rD)
se_na <- SummarizedExperiment(assays = list(jiang2019_ibaq_na),
    colData = cD, rowData = rD)
```

After creating a `SummarizedExperiment` object, the shiny application can be started via

```
shinyQC(se)
```

The `MatrixQCvis` package offers several visualizations to explore the data quality and underlying statistical properties of the data set. Some of these visualizations will be plotted in the following paragraphs.

#### 2.1 Normalization, transformation and missing value imputation of the data set of Jiang et al. (2019)

MaxQuant assigns the value 0 to the intensity table if there was not enough information to quantify the peptide/protein. From their Methods section it is not clear how Jiang et al. [2019] dealt with missing values, in particular, whether

1. they continued with zero-imputed values or
2. they set intensity values with 0s to NA and imputed later by a more sophisticated method (cf. Lazar et al. [2016]).

The aim of a normalization strategy is to reduce the effects of technical variability on the data set. Jiang et al. [2019] performed quantile normalization in their analysis.

Normalization of data is complicated and there is no universal, *a priori* normalization strategy available for different data sets. Rather, the normalization technique has to be chosen based on the properties of the data set and the choice is likely to produce erroneous results when the underlying assumptions are not met [Wang et al., 2011, Wu et al., 2014]. In the case of quantile normalization, it is assumed that all samples have the same distribution regardless of the sample type. This assumption, however, is only valid when a small number of proteins is disregulated [Wu et al., 2014]. Violations of these assumptions may lead to false positive results, effect-size reduction, and masking of true effects; more nuanced methods, such as class-specified normalization, in which the data is split based on the sample class/type, after which quantile normalization is performed on each split separately, might be more suitable in these cases [Zhao et al., 2020].

Using the naive quantile normalization approach and the MaxQuant data set from Jiang et al. [2019] containing 0s distorted the distributions of the underlying data (see Figures 1 and 2, `limma::normalizeQuantiles` used as in Jiang et al. [2019]). On the other hand, performing quantile normalization on the data set containing NA values, resulted in identical distribution of the samples. From the Methods section in Jiang et al. [2019] it became not clear if the quantile normalization was done on the data set containing 0s or NAs; Extended Data Fig. 2 in Jiang et al. [2019] shows after $\log_2$ transformation non-identical distributions, contrary to the expected result from quantile normalization (see Figure 1 d).

We argue that visualization of the normalization results, as offered by `MatrixQCvis`, e.g. by looking at the per-sample distribution before and after conducting normalization or visualizations of trend lines of the per-sample median values or sum of values, will provide valuable insight into the expected execution of normalization strategies (i.e. reduction of technical variability and distribution properties of the underlying data).

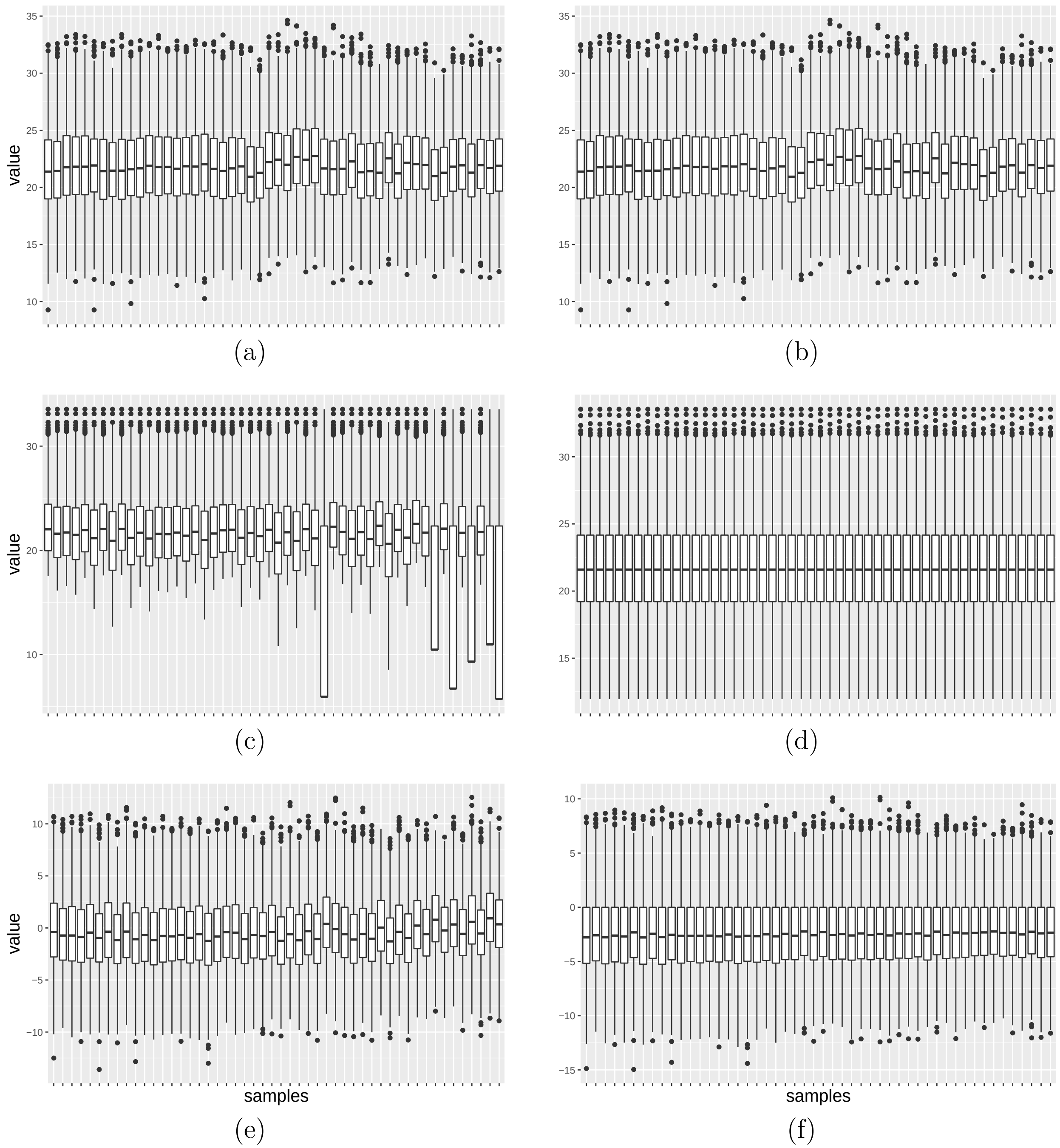


Figure 1: Sample-wise boxplots of intensity values. (a) raw values (containing missing values as 0), (b) raw values (containing missing values as NA), (c) normalized values (quantile normalization, missing values as 0), (d) normalized values (quantile normalization, missing values as NA), (e) normalized values (75% quantile division, missing values as 0), (f) normalized values (75% quantile division, missing values as NA). All values were $\log_2$-transformed prior to visualization. To improve readability only a subset of 50 samples is shown.

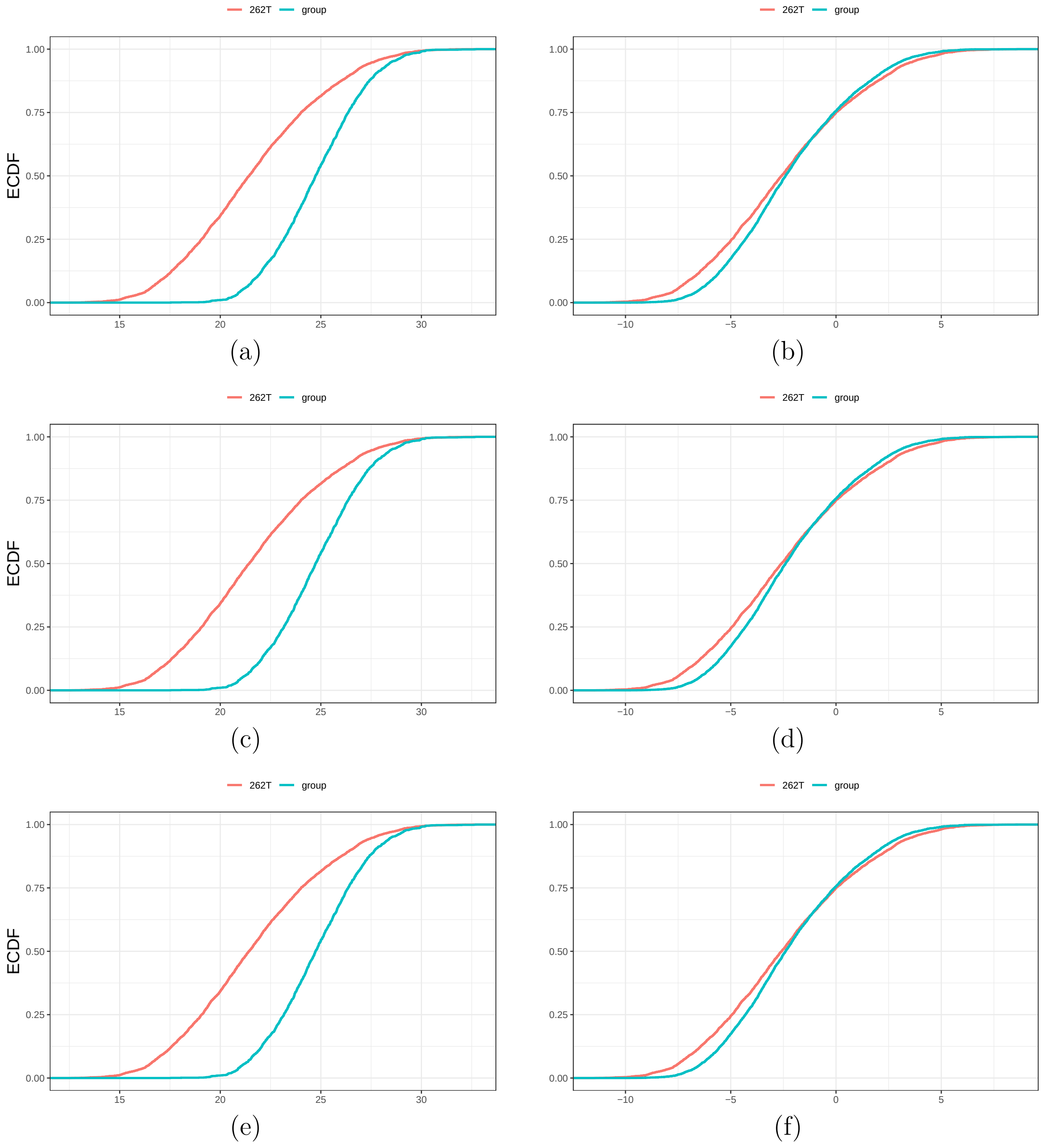


Figure 2: Empirical cumulative distribution function (ECDF) plots of intensity values. Shown are the ECDF plots of the sample 262T and all the means of the samples belonging to the tumor samples (group) except sample 262T. (a) raw values (containing missing values as 0), (b) raw values (containing missing values as NA), (c) normalized values (quantile normalization, missing values as 0), (d) normalized values (quantile normalization, missing values as NA), (e) normalized values (75% quantile division, missing values as 0), (f) normalized values (75% quantile division, missing values as NA). All values were $\log_2$-transformed prior to visualization.

#### 2.2 Missing values in the data set of Jiang et al. (2019)

When detecting missing values in the `SummarizedExperiment` object, `MatrixQCvis` will load a dedicated interface to analyze missing and measured values. Missing values are an inherent characteristic of proteomics and metabolomics data sets acquired by mass spectrometry. Poor sample quality, however, might be characterized by a higher number of missing values and different sample types might show a different distribution of missing values. `MatricQCvis` offers visualizations that help to analyze these relations.

In addition, proper handling of missing values is required for sound biological information inference. `MatrixQCvis` includes several imputation methods [Lazar et al., 2016] for missing values; furthermore, `MatrixQCvis` enables to perform differential testing on the non-imputed data sets using the `proDA` package [Ahlmann-Eltze and Anders, 2019].

Figure 3 will showcase some of the included capabilities to display and analyze the distribution of measured values.

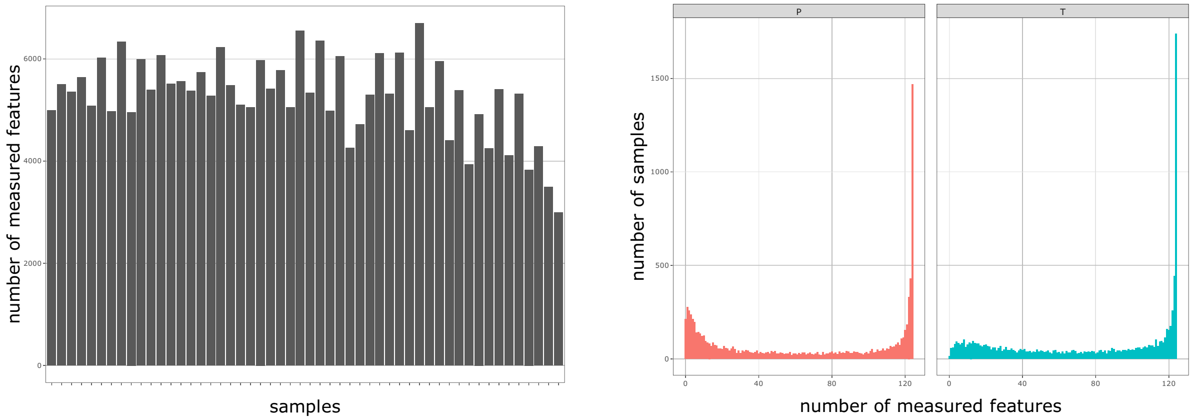


(a) (b)

Figure 3: Measured values in proteomics data set of Jiang et al. [2019]. (a) Barplot displaying the number of measured values along samples. The samples XL21P and XL18P have with 2992 and 3497 measured features the lowest and second lowest number of features in the data set. In total there are 9251 features in the data set. For better readibility only the first 50 samples and the samples XL21P and XL18P are shown. (b) Histogram of number of features along all samples stratified for the tissue type, non-tumor paired (P) and tumor (T) tissue. Shown are the number of features (y-axis) with a given number of measured values per feature along the feature (x-axis).

#### 2.3 Trends/drifts in the data set of Jiang et al. (2019)

A diagnostic plot to identify trends in the data set stemming from technical variation is the trend/drift plot that shows the ordered set of samples on the x-axis and aggregated information (median or sum of values per sample) on the y-axis. This plot is valuable to identify systematic trends in the data set (e.g. due to fluctuations in mass spectrometry sensitivity dependent on acquisition time or batch effects). The trend lines should be checked after application of normalization, transformation, batch correction, and imputation methods.

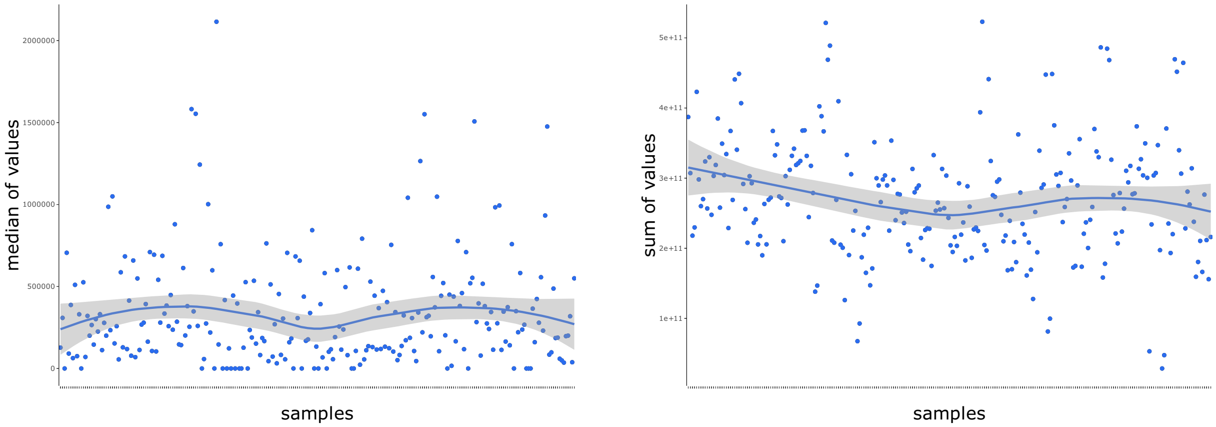


(a) (b)

Figure 4: Trend/drift plots of intensity values. Each point represents a sample. (a) Intensity values are aggregated by the median of all values of a sample, (b) intensity values are aggregated by the sum of all values of a sample. The indicated trend line (by LOESS smoothing) serves rather as a visual help to indicate constant drifts in the data set and might not be appropriate if there are abrupt effects (i.e. a step in the median or sum of intensities).

#### 2.4 Analysis of homogeneity of variance of the data set of Jiang et al. (2019)

Mean vs. standard deviation plots (mean-sd plots) display the (ranked) means of the features on the x-axis and the standard deviation of the features on the y-axis (see Figure 5). The combination of normalization, transformation, batch correction and imputation should result in a data set where the standard deviation is independent of the mean, i.e. that the features have the same finite variance (homogeneity of variance). The red lines in the plots indicate the running median estimator (window width 10%), and, in the case of mean-standard deviation independence, the running median estimator should be approximately horizontal. Imputation leads to the distortion of homogeneity of variance (see Figure 5 c and d). The effect has to be taken into account when performing differential expression analysis by including information on variance of the probed features.

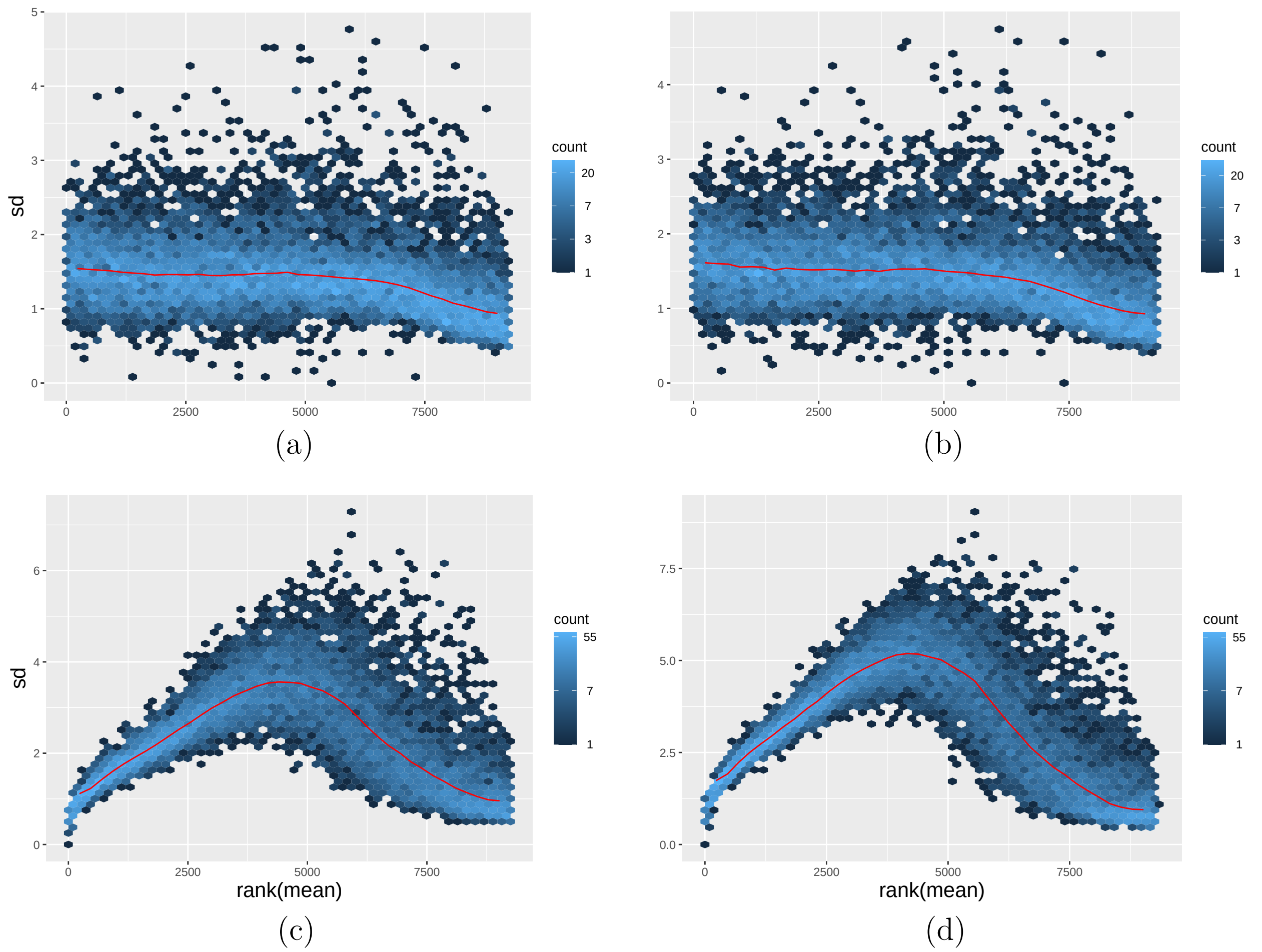


Figure 5: Mean vs. standard deviation plot. (a) transformed values (quantile normalization), (b) transformed values (75% quantile division normalization), (c) imputed values (quantile normalization), (d) imputed values (75% quantile division normalization).

#### 2.5 Intensity values per feature of the data set of Jiang et al. (2019)

The shiny application enables to display the values stored in the `assay` slot of the `SummarizedExperiment` object. In the case of the proteomics data set of Jiang et al. [2019], these correspond to intensity values. The values are available via the sub-tab `Features` in the tab `Values`. The panel shows the values per sample of the raw values and after the processing steps normalization, transformation, batch correction and imputation (see Figure 6 a).

The panel also displays another figure showing the coefficients of variation for the features per data processing step, see Figure 6 b). This plot might prove useful when identifying features that show high variation of across the samples, e.g. due to technical or experimental effects.

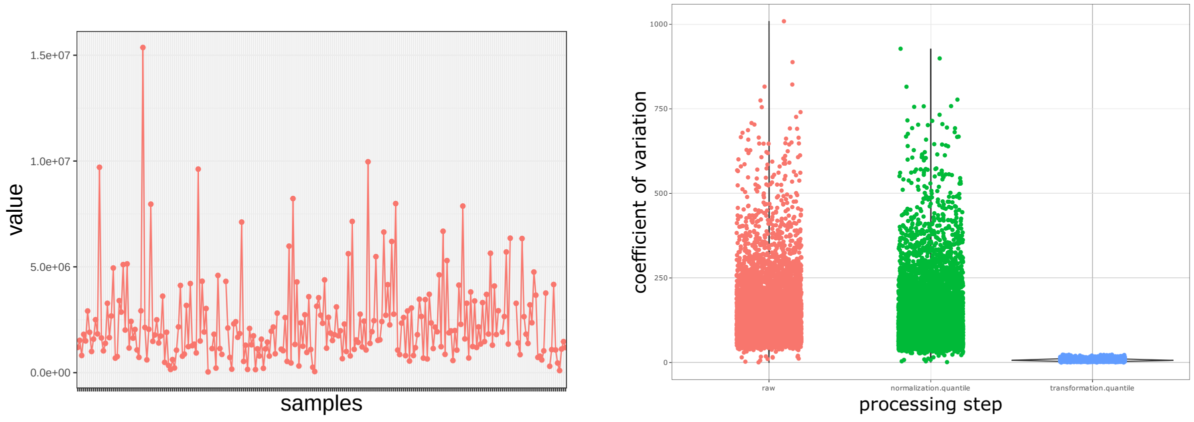


(a) (b)

Figure 6: Intensity values. (a) normalized intensity values (quantile normalization) across samples for the feature A0AV96 (RNA-binding protein 47), (b) coefficient of variation for features for the raw, normalized (quantile normalization), and $\log_2$ transformed values.

#### 2.6 Dimension reduction of the data set of Jiang et al. (2019)

`MatrixQCvis` includes functions to create dimension reduction plots (principal component analysis, PCA; principal coordinates analysis/ multidimensional scaling, PCoA/MDS; non-metric multidimensional scaling, NMDS; t-distributed stochastic neighbor embedding, tSNE, van der Maaten and Hinton [2008]; uniform manifold approximation and projection, UMAP, McInnes et al. [2018]). Dimension reduction is a technique to show and explore the relations between variables of the underlying data in a reduced dimension space. All dimension reduction techniques employ geometrical projections that take points from a high- into a low-dimensional space. Next to the linear techniques (PCA, PCoA), non-linear dimension reduction techniques (NMDS, tSNE, UMAP) are implemented in the `MatrixQCvis` package.

As an example, PCA will reduce the 248-dimensional space from the data set of Jiang et al. [2019] (consisting of 248 sample points) to a 2-dimensional plane (principal components 1 and 2) that explain 16.9% and 4.9%, respectively, of the variance of the 248 dimensional data set. `MatrixQCvis` enables to highlight the different factors of columns in `colData(se)` by different colors (see Figure 7).

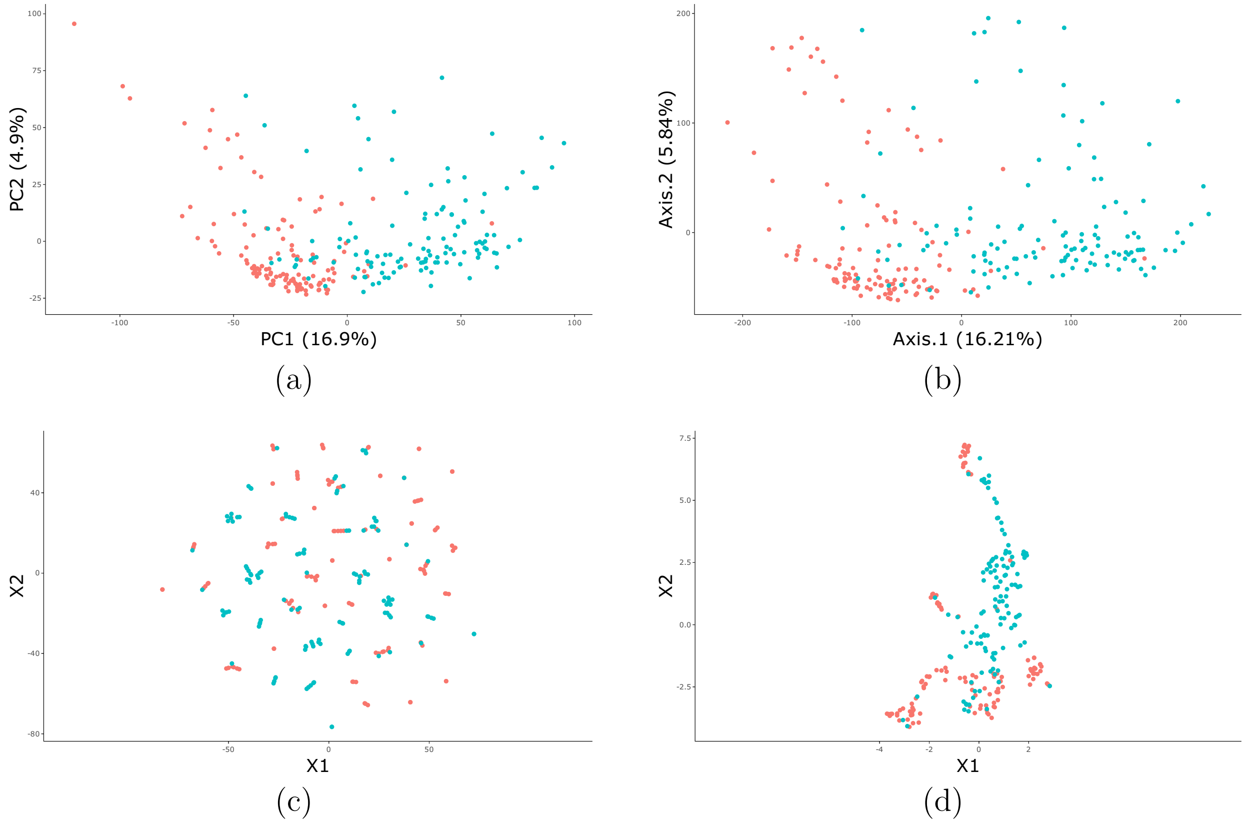


Figure 7: Dimension reduction plots for the data set of Jiang et al. [2019] (quantile normalization, $\log_2$ transformation, MinDet imputation). (a) principal component analysis, (b) principal coordinates analysis/multidimensional scaling, (c) t-distributed stochastic neighbor embedding, (d) uniform manifold approximation and projection. Features with a standard deviation of 0 were removed prior to performing dimension reduction. The red-colored features correspond to the non-tumor paired tissue samples, the blue-colored features correspond to tumor samples.

#### 2.7 Differential expression analysis of the data set of Jiang et al. (2019)

Using the `MatrixQCvis` package, it is possible to recapitulate the differential expression analysis of Jiang et al. [2019] (see Supplementary Table 6 therein). Before conducting the expression analysis, the (paired) samples that showed an increased number of missing proteins will be removed prior to conducting the differential expression analysis (this will be the samples XL21P, XL21T, XL18P, and XL18T). To conduct a differential expression analysis of paired samples, the following design formula will be used:

$$\sim \ 0 + patient + cell\_type \tag{1}$$

where *patient* is the column in the `colData` of the `SummarizedExperiment` that contains the identifier of a patient and *cell_type* will be either `T` or `P` depending if the proteins were measured from tumor (`T`) or paired non-tumor (P) samples.

The specified contrast will be *"cell_typeT"*, i.e. the effect between the *cell_type* `T` and `P` will be tested.

`MatrixQCvis` will display the result of a differential expression analysis both in a tabular format (similar to Table 1) and in the form of a volcano plot (see Figure 8). The employed normalization, transformation and imputation strategies and exclusion of the above-mentioned sample pairs due to increased number of missing values resulted in comparable results to Jiang et al. [2019] based on moderated t-tests (see Table 1).

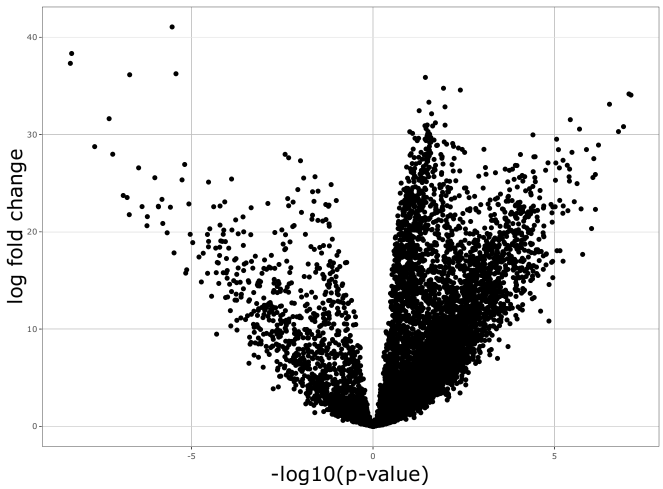


Figure 8: Volcano plot. Each point represents a protein. The shiny application will create an interactive figure of the volcano plot that displays the name, the adjusted p-value, the -$\log_{10}$(adjusted p-value), and the log fold change upon hovering over the protein. Moderated t-tests were performed with Benjamini-Hochberg adjustment for multiple testing.

Table 1: Differential expression between tumor and non-tumor paired tissue samples for the top 20 differentially expressed proteins. The underlying data set contains the 9251 features from Jiang et al. (2019). Within Jiang et al. (2019) only a subset of 2008 features was reported in the differential expression analysis (Supplementary Table 6 in Jiang et al., 2019). The analysis is based on performing moderated t-tests. The features are ordered according to the Benjamini-Hochberg adjusted p-values. Shown are the log fold changes (logFC) from our analysis and Jiang et al. (2019) and the corresponding ranks of the features based on Benjamini-Hochberg adjusted p-values from Jiang et al. (2019).

| Uniprot_ID | logFC (limma) | logFC (Jiang et al., 2019) | rank (Jiang et al. 2019) |
|---|---:|---:|---:|
| Q8WWQ8 | -5.53 | -5.67 | 9 |
| Q6UXB4 | -8.30 | -9.32 | 1 |
| Q9H2X3 | -8.34 | -8.75 | 4 |
| Q8WWZ8 | -5.43 | -5.57 | 32 |
| Q9UEW3 | -6.70 | -7.19 | 6 |
| P08238 | 1.44 | | |
| P78527 | 1.94 | 2.02 | 5 |
| P02751 | 2.41 | 2.54 | 14 |
| Q9BX97 | 7.05 | 7.82 | 2 |
| P52292 | 7.11 | 7.76 | 3 |
| P34932 | 1.54 | | |
| P55290 | 6.51 | 6.91 | 7 |
| Q6PI48 | 1.98 | 2.17 | 16 |
| P54578 | 1.27 | | |
| O43776 | 1.61 | | |
| Q9BYV7 | -7.27 | -8.18 | 13 |
| Q16363 | 5.43 | 5.80 | 19 |
| Q9H223 | 1.72 | | |
| Q9UMS4 | 1.49 | | |
| Q9H3P7 | 1.98 | 2.05 | 8 |

### 3 Brueffer et al. (2018): Clinical Value of RNA Sequencing-Based Classifiers for Prediction of the Five Conventional Breast Cancer Biomarkers

In a second application, we will highlight the visualization functions of `MatrixQCvis` on a RNA-sequencing data set from breast cancer biopsies.

The data set stems from the population-based multicenter Sweden Cancerome Analysis Network - Breast Initiative [Brueffer et al., 2018, 2020] and included Illumina paired-end RNA-sequencing data from 405 patients. The data set was accessed via the GEO database (accession number GSE81538). The data set contains comprehensive multi-rater histopathologic evaluation and was used as a training set for a classifier for a larger set consisting of 3273 samples (accession number GSE96058).

In a first step, the data set is loaded and converted to a `SummarizedExperiment` object that is readable by the `shinyQC` function. Furthermore, the `SummarizedExperiment` will be truncated in this analysis here: FPKM (fragments per kilobase of transcript per million mapped reads)-normalized transcripts with a mean expression below zero and invariant features (with standard deviation below 0.5) will be removed.

```
f <- "Brueffer_et_al_2018/GSE81538_gene_expression_405_transformed.csv"
brueffer2018 <- read.table(f, sep = ",", header = TRUE,
    row.names = 1)

## assay
a <- as.matrix(brueffer2018)

## colData
cD <- DataFrame(name = colnames(a))

## rowData
rD <- DataFrame(features = rownames(a))

## create the SummarizedExperiment object
se <- SummarizedExperiment(assays = a, colData = cD, rowData = rD)

## filter the transcripts based on mean expression and standard deviation
se <- se[apply(a, 1, mean) >= 0 & apply(a, 1, sd) >= 0.5, ]
```

After creating the `SummarizedExperiment` object the shiny application can be started via

```
shinyQC(se)
```

#### 3.1 Transformation of the data set of Brueffer et al. (2018) and analysis of homogeneity of variance

The expression values are already FPKM-normalized. Here, the FPKM values will be subjected to variance stabilizing normalization/transformation only.

To check the statistical properties and the effects of variance stabilizing transformation, mean vs. standard deviation plots will give information on the homogeneity of variance across the mean-ranked features. Ideally, these plots show an approximately horizontally running median estimator (red line, window width 10%, cf. Figure 9).

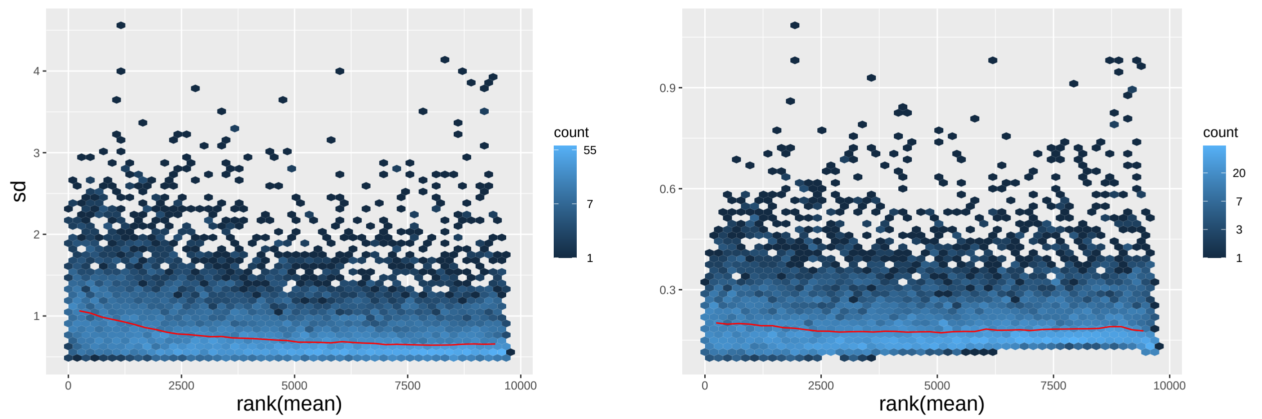


(a) (b)

Figure 9: Mean vs. standard deviation plot for (a) FPKM-normalized and (b) transformed data (variance stabilization normalization/transformation) of the data set of Brueffer et al. [2018]. The normalized and transformed transcript values show variance homogeneity.

#### 3.2 MA plots and Hoeffding's $D$ statistic values for the data set of Brueffer et al. (2018)

Next to the visualization plots shown above for the proteomics data set of Jiang et al. [2019], `MatrixQCvis` contains also other visualizations, such as MA plots (see Figure 10), to assess the data quality. Commonly, MA plots are used to identify abnormalities in the sample. The MA plot facilitates to identify samples that show abnormal values compared to a group of other samples (e.g. other samples of disease/healthy type).

Here, $M$ is defined as

$$M = I_i - I_j \tag{2}$$

and $A$ is defined as

$$A = \frac{1}{2} \cdot (I_i + I_j) \tag{3}$$

where $I_i$ and $I_j$ are the logarithms of intensity or count values. The values for $I_i$ are taken from the sample $i$. For $I_j$, the feature-wise means are calculated from the values of the group $j$ of samples. The sample for calculating $I_i$ is excluded from the group $j$.

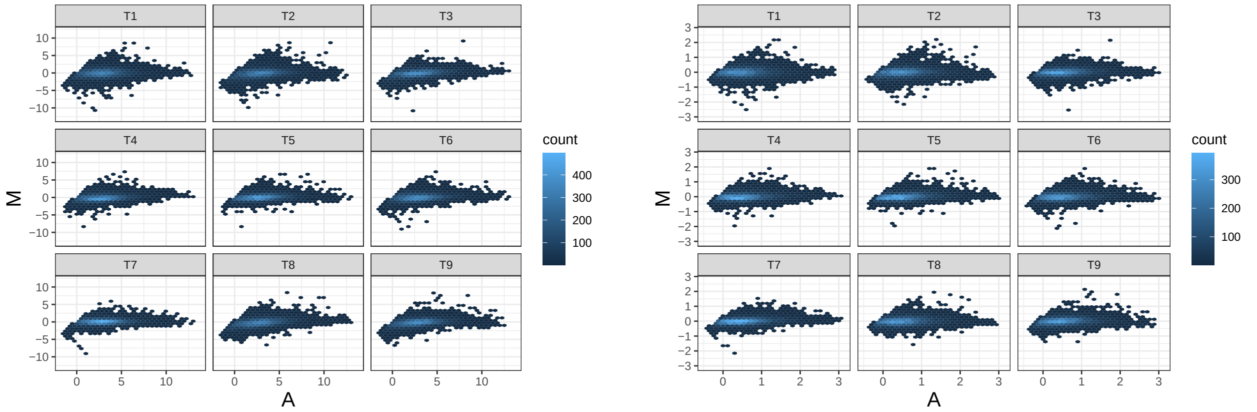


(a) (b)

Figure 10: MA plots of the first nine samples, T1-T9, for the (a) FPKM-normalized and (b) transformed values (variance stabilization normalization/transformation). The group consists of the per-feature mean values of all samples expect the respectively displayed sample (T1, ..., T9). The MA plot does not indicate that any sample in the data set of Brueffer et al. [2018] is an outlier.

The MA plots can be summarized in the Hoeffding's $D$ statistic plot where each point refers to a sample in the data set (see Figure 11). $D$ is a measure of the distance between $F(A, M)$ and $G(A)H(M)$, where $F(A, M)$ is the joint cumulative distribution function (CDF) of $A$ and $M$, and $G$ and $H$ are marginal CDFs. The higher the value of $D$, the more dependent are $A$ and $M$.

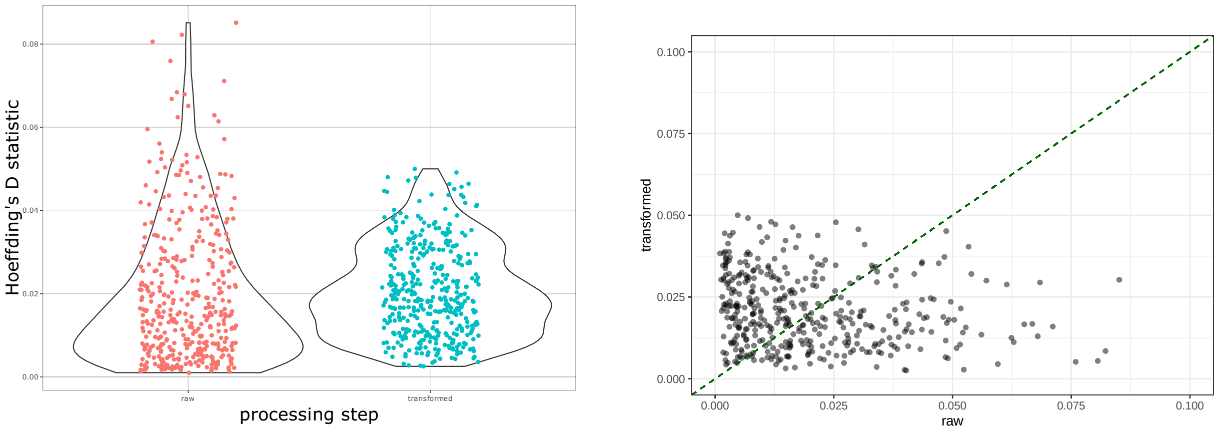


(a) (b)

Figure 11: Hoeffding's $D$ statistic values. (a) Hoeffding's $D$ statistic plot for the raw and transformed (variance stabilization) values. Each dot represents a sample. Transformation increases overall Hoeffding's D statistic values, but reduces the D statistic values of samples after normalization with extreme D statistic values. In general, Hoeffding's D statistic values are only moderately high for some of the samples, indicating that there is no indication for outlier samples from this diagnostic plot. (b) Scatterplot between Hoeffding's $D$ statistic values of raw and transformed values. The identity line is shown as a green dashed line.

#### 3.3 Additional diagnostic plots to detect outlier samples in the data set of Brueffer et al. (2018)

Other diagnostic plots in exploratory data analysis to identify outlier samples are empirical cumulative distribution function (ECDF) plots (see Figure 12). The ECDF gives information on the distribution properties of a single sample and enables to identify the proportion of values below (or above) a certain threshold. `MatrixQCvis` includes functionality to calculate an averaged ECDF based on the specified group. Comparing the ECDF of a sample with the ECDF of the specified group, enables to add another layer of information if a sample deviates from the expected distribution (based on the group properties).

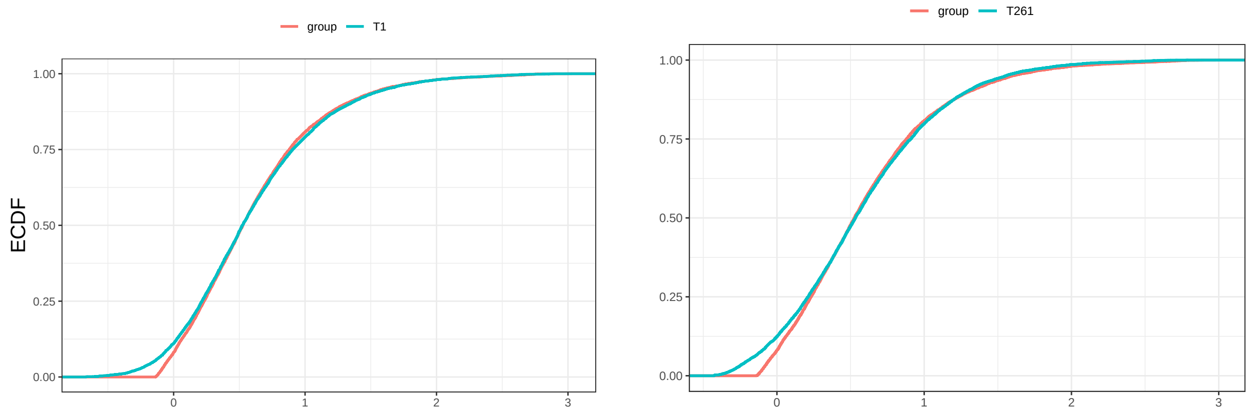


(a) (b)

Figure 12: Empirical cumulative distribution function (ECDF) plots of samples (a) T1 and (b) T261 after variance stabilization. The 'group' consists of the per-feature mean values of all samples expect the selected one (T1, T261).

Another diagnostic plot is based on distances between samples. `MatrixQCvis` enables to calculate distances between samples and provides visualizations to display these. Outlier samples might form (singleton) clusters within the heatmap, i.e. they might have a higher distance to other samples, and will show higher sums of distances.

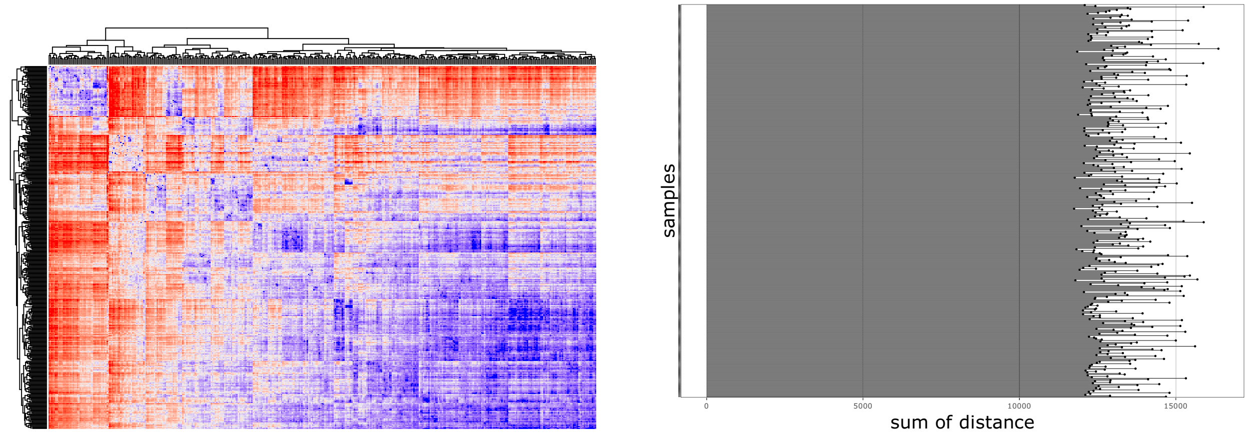


(a) (b)

Figure 13: Distances between samples (after transformation). (a) Heatmap of euclidean distances. (b) Sum of euclidean distances of one sample to other samples. The plots show that there are no samples that deviate substantially from the set of others (no singleton clusters with high distances, no samples with large sum of distance to other samples).

Taken together, the QC analysis conducted here using the `MatrixQCvis` package suggested that there is no indication that the data set of Brueffer et al. [2018] contains any outlier samples and that the data set shows favored statistical properties after FPKM-normalization and variance-stabilizing normalization/transformation (e.g. variance homogeneity across the features).
